# Characterizing transcriptional regulatory sequences in coronaviruses and their role in recombination

**DOI:** 10.1101/2020.06.21.163410

**Authors:** Yiyan Yang, Wei Yan, A. Brantley Hall, Xiaofang Jiang

## Abstract

Novel coronaviruses, including SARS-CoV-2, SARS, and MERS, often originate from recombination events. The mechanism of recombination in RNA viruses is template switching. Coronavirus transcription also involves template switching at specific regions, called transcriptional regulatory sequences (TRS). It is hypothesized but not yet verified that TRS sites are prone to recombination events. Here, we developed a tool called SuPER to systematically identify TRS in coronavirus genomes and then investigated whether recombination is more common at TRS. We ran SuPER on 506 coronavirus genomes and identified 465 TRS-L and 3509 TRS-B. We found that the TRS-L core sequence (CS) and the secondary structure of the leader sequence are generally conserved within coronavirus genera but different between genera. By examining the location of recombination breakpoints with respect to TRS-B CS, we observed that recombination hotspots are more frequently co-located with TRS-B sites than expected.

## INTRODUCTION

In the past two decades, at least three novel coronaviruses have spilled over from animals into humans, SARS, MERS, and SARS-CoV-2(Cui et al., 2019; Guarner, 2020). Coronaviruses are positive-sense, single-stranded RNA viruses with large genomes and are grouped into four genera: Alphacoronaviruses, Betacoronaviruses, Gammacoronaviruses, and Deltacoronaviruses(Cui et al., 2019). Recombination between coronaviruses plays an important role in coronavirus evolution and can alter host range, pathogenicity, and transmission pattern(Bentley and Evans, 2018; Graham et al., 2018; Keck et al., 1988; Lai et al., 1985; Lau et al., 2010; Tian et al., 2014; Wang et al., 2017, 1993; Xiao et al., 2017; Zhang et al., 2005). Inter-coronavirus recombination requires two different but related coronaviruses to co-infect a cell(Graham et al., 2018; Graham and Baric, 2010; Simon-Loriere and Holmes, 2011). Recombination in coronaviruses occurs during genome replication when the RNA-dependent RNA polymerase (RdRp) replicating the genome dissociates from one viral genome currently serving as the template and reassociates with a different viral genome while retaining the nascent RNA in a process called template switching(Sawicki and Sawicki, 1998; Simon-Loriere and Holmes, 2011). Therefore, template switching generates a recombinant RNA originating from the genomes of two coronaviruses(Bentley and Evans, 2018; Simon-Loriere and Holmes, 2011).

Coronavirus and transcription also involves template switching(Sawicki and Sawicki, 1998). After a coronavirus infects a cell, it replicates its positive-strand RNA genome into a negative strand genome with RdRp(Sawicki and Sawicki, 1998). The negative strand genome subsequently serves as a template for the production of positive-strand genomes and subgenomic messenger RNAs (sgmRNAs), a set of 3’ coterminal RNAs encoding structural genes(Sawicki and Sawicki, 1998). The sgmRNAs share a common 5’ sequence, called a leader sequence, which is located at the beginning of the coronavirus genome(Zhang et al., 1994). The leader sequence is added to the 5’ end of all sgmRNAs through RdRp template switching(Sawicki and Sawicki, 2005, 1998). Template switching occurs as RdRp is transcribing the negative strand and encounters transcriptional regulatory sequences (TRS) preceding each gene called the body TRS (TRS-B)(Sola et al., 2015). The TRS-B site has a 7-8 bp conserved core sequence (CS) which is thought to enhance the likelihood of RdRp template switching by hybridizing with an identical or nearly identical CS in the leader TRS (TRS-L)(Sola et al., 2015; Zúñiga et al., 2004). The occurrence of this programmed template switching leads to the generation of sgmRNAs with identical 5’ and 3’ sequences, but alternative central regions corresponding to the beginning of each structural ORF(Sawicki et al., 2007; Sawicki and Sawicki, 2005, 1998; Sola et al., 2015; Wu and Brian, 2007).

Because TRS-B is a signal for RdRp to switch templates, it is reasonable to hypothesize that recombination events are more likely to occur at or near TRS-B sites(Graham et al., 2018). Once RdRp has dissociated from the original template after encountering TRS-B, it could reassociate with a different genome, leading to recombination. Identical or nearly identical TRS sites between different coronaviruses could hybridize promoting recombination at TRS-B sites. Therefore, in this study we investigate the relationship between genome recombination in coronaviruses and template switching at TRS-B sites. TRS-B sites are not annotated in every coronavirus genome because there is not a systematic tool to identify them nor has there been a project focused on TRS-B annotation. Therefore, there has not been a systematic study of whether recombination events are more common around TRS-B sites. Thus, we developed a tool called SuPER to systematically identify TRS sites within coronavirus genomes to better understand the relationship between coronavirus recombination and template switching. After systematically identifying TRS sites and recombination events in coronaviruses, we found that 8 of 91 (8.7%) of TRS-B sites are within breakpoint hotspots and that recombination hotspots are more frequently co-located with TRS-B sites than expected.

## RESULTS

### Systematic identification of TRS in coronavirus genomes

To examine how template switching at TRS-B sites contributes to genome recombination in coronaviruses, we needed to systematically identify TRS-B sites. Therefore, we developed the tool SuPER to identify TRS-B sites in coronavirus genomes (Figure 1A). SuPER first uses a covariance model derived from Rfam to identify TRS-L via profile-based sequence and structure scoring. Then, SuPER identifies TRS-B sites either by identifying template switching junctions with RNA-seq or in the absence of RNA-seq by identifying sequences preceding genes that are similar to the TRS-L CS as putative TRS-B CS.

**Figure 1:**
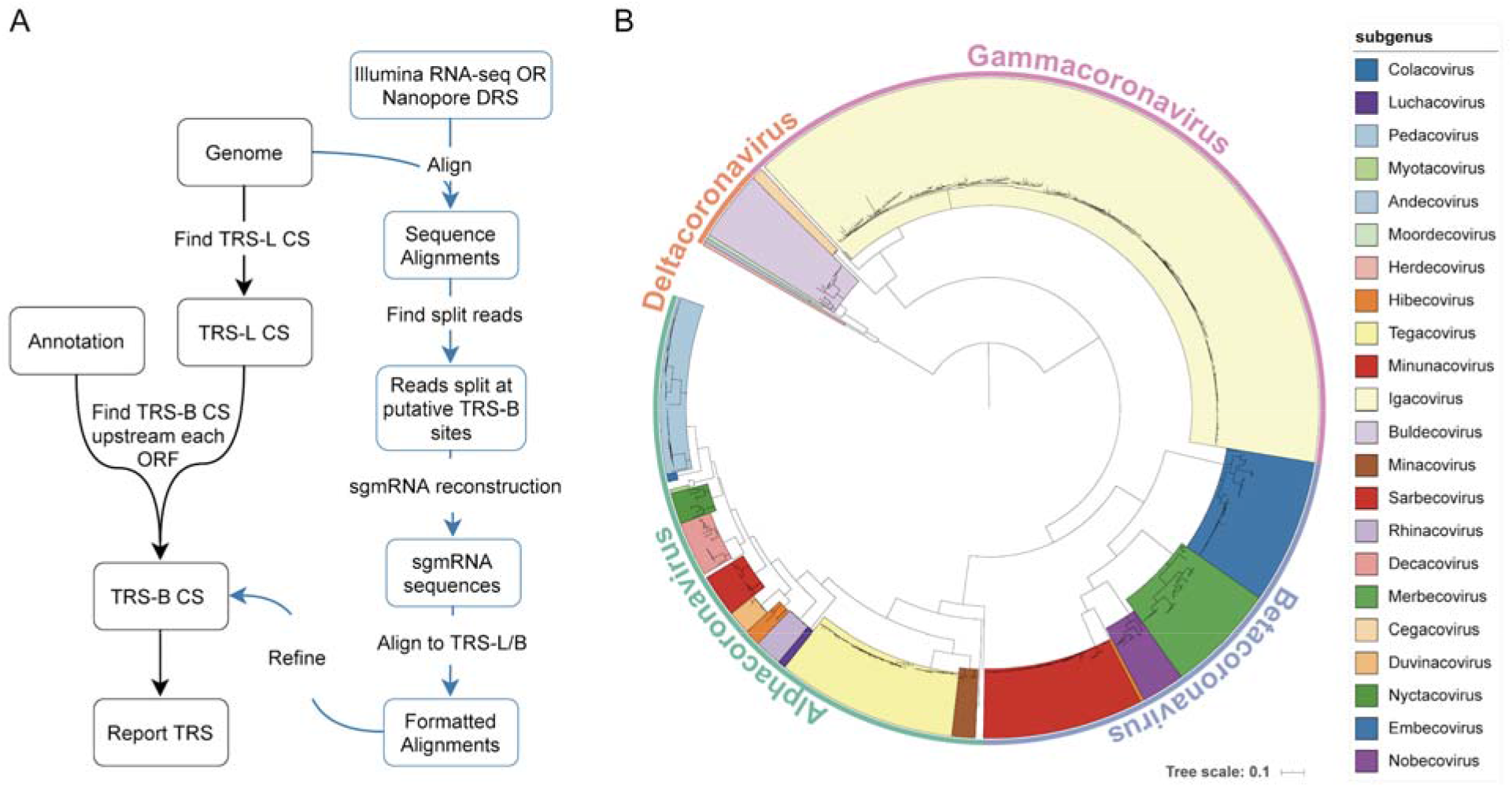
**A.** SuPER workflow. The main pipeline is shown with black text boxes and arrows. Analysis procedures specific to sequencing data are shown with blue text and arrows. **B.** The phylogenetic tree built from RdRp from the 506 representative coronavirus genomes used in the study.

To validate SuPER, we ran it on the SARS and SARS-CoV-2 genomes. The TRS-L/B of SARS and SARS-CoV-2 can be identified or inferred from previous research (Kim et al., 2020; Rota, 2003). We found that when using SuPER with RNA-seq data, we can correctly identify all known TRS-B sites from these two genomes with zero false positives (STable 1). When running SuPER without RNA-seq data, if only high confidence TRS-B sites are reported, SuPER can predict all known 16 TRS-B CS accurately and precisely. SuPER also reports six extra TRS-B but labeles them as “not recommended” due to the fact that their hamming distance to TRS-L CS is more than 1 base pair. The reason we still report low confidence TRS-B is due to the concern of missing potential non-canonical TRS-B CS.

To systematically identify TRS-B in coronaviruses, we used a set of 506 non-redundant representative genomes from the family Coronaviridae (STable1; See methods). Not all coronavirus genomes were assigned to a subgenus in the NCBI Virus database. To place the unassigned genomes into genera and subgenera, we built a phylogenetic tree based on the RdRp protein sequence(Wolf et al., 2018) from all representative genomes (Figure 1B). In total, we assigned 498 genomes into 23 subgenera (STable 2). We identified putative TRS-L and TRS-B CS in the 506 coronavirus genomes with SuPER. Of those, SuPER was run on 11 genomes with RNA-seq data and the remaining genomes without RNA-seq data (STable 3 and STable 4). In total, SuPER identified 465 TRS-L and 3509 TRS-B.

### TRS characterization in coronavirus genomes

Using the results from SuPER, we examined the conservation of TRS-L CS and its position in the secondary structure in all coronavirus subgenera (Figure 2A). In general, we found that the TRS-L CS is conserved within genera but differs between genera, with the exception of Embecoviruses where the TRS-L CS is similar to Alphacoronaviruses. The secondary structure of the leader sequences were visualized by VARNA(Darty et al., 2009) (SFigure 1). We observed that the secondary structure of the leader sequence is relatively conserved within subgenera, but different between subgenera.

**Figure 2:**
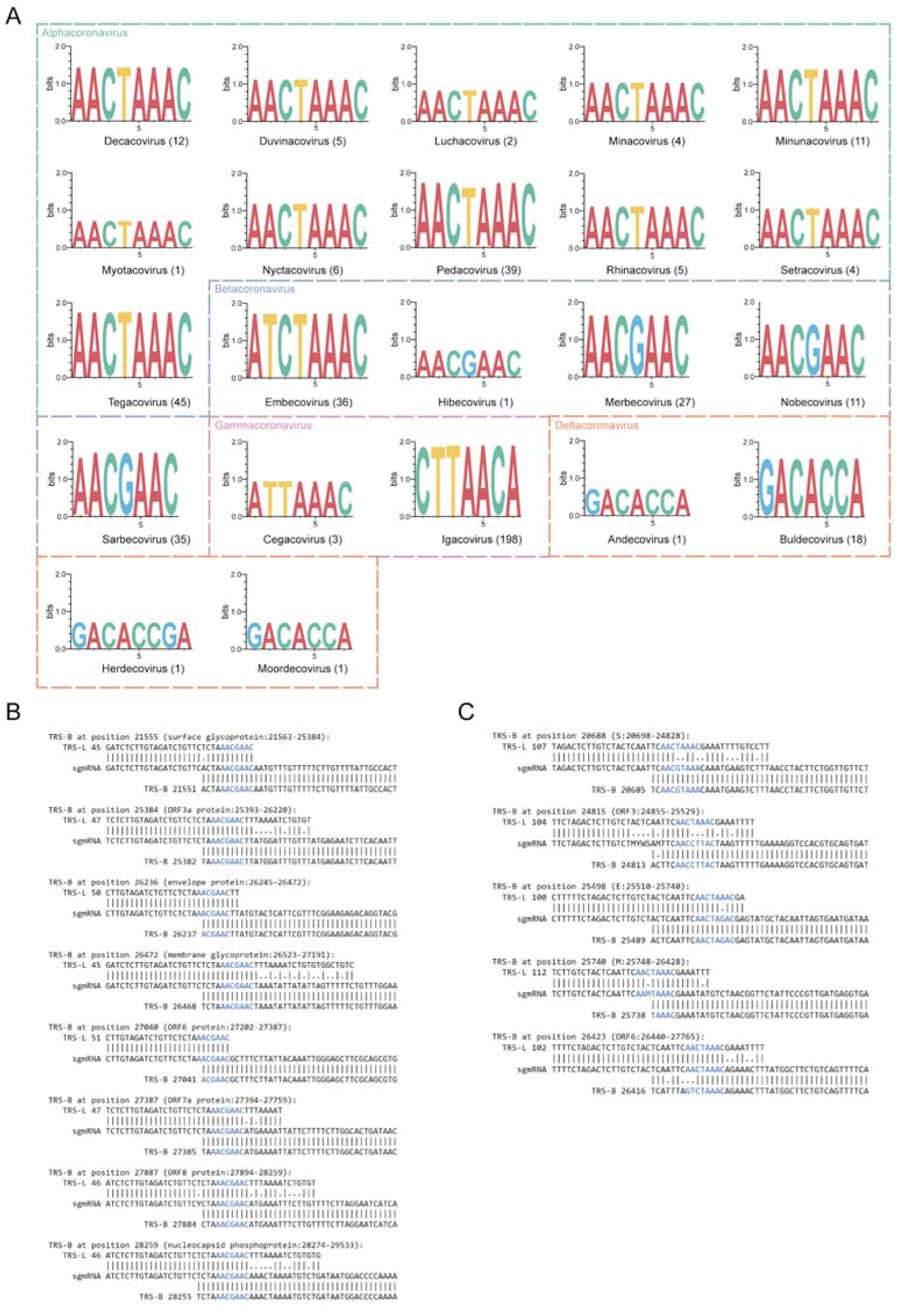
**A.** Sequence logo of the TRS-L CS from all coronavirus subgenera.The number following the subgenus name indicates the number of sequences used to generate the sequence logo. **B** and **C.** Alignments of TRS-L, sgmRNA, and TRS-B to illustrate the template switching pattern in SARS-CoV-2. (B) and Porcine epidemic diarrhea virus (C). CS in TRS-B, TRS-L and sgmRNA are colored blue.

We examined the TRS-B CS detected from coronavirus genomes where RNA-seq was available using SuPER. RNA-seq data can help to identify the template switching patterns and accurately detect the TRS-B CS. We found that TRS-B CS were either identical to TRS-L CS from the same genome, or differed by only one base pair in most genomes including SARS-CoV-2, SARS, MERS, SADS-CoV and HCoV-HKU1 (Figure 2B). However, in some cases, there is limited similarity between TRS-L and TRS-B, making it difficult to identify TRS-B without RNA-seq data. For example, in Porcine Epidemic Diarrhea Virus, the TRS-B CS can differ by more than two base pairs from the TRS-L CS (Figure 2C). We also observed similarity in a few base pairs upstream and downstream of the CS, which could be critical for mediating base-pairing during template switching in cases where the TRS-L and TRS-B CS differ (Figure 2B)(Sola et al., 2005). Not all annotated ORFs preceded by TRS-B sites are supported by RNA-seq data. This could be due to the lack of enough coverage to detect the template switching events occurring at that position or there is low or no template switching. For example, we did not find RNA-seq data supporting the TRS-B site proceeding ORF7b and ORF10 with SuPER, and a previous study using Oxford Nanopore direct RNA sequencing was unable to validate the existence of a sgmRNA corresponding to ORF7b and ORF10(Kim et al., 2020).

### Detecting recombination in coronaviruses

We performed recombination analysis on the genomes for each subgenus using RDP4(Martin et al., 2015). We chose the subgenus level due to the fact that genomes from different subgenera often share less than 50% nucleotide identity leading to poor alignments, causing issues for recombination detection. By building phylogenetic trees from RdRp and the structural genes S, E, M, and N from Betacoronaviruses, we found that each subgenus formed a distinct clade for RdRp and the structural genes, providing no evidence of recent inter-subgenus recombination (Figure 3A). On the contrary, we found conflicting branching orders and incongruent topology of the subgenera phylogenies suggesting recombination events within subgenera. In total, we detected 973 recombination events in 16 subgenera with RDP4 (SData 1). The number of recombination events detected is largely dependent on the number and diversity of representative genomes available for each subgenus.

**Figure 3:**
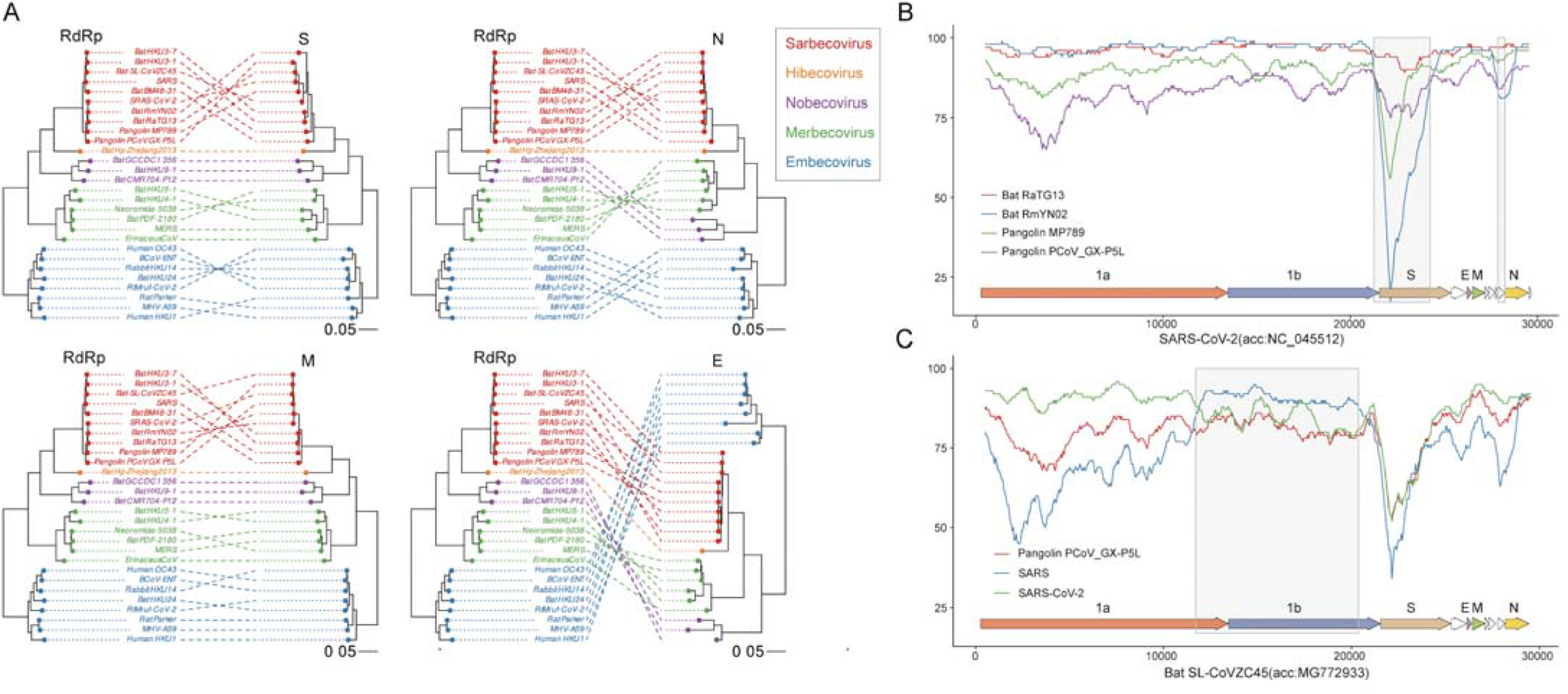
**A.** Tanglegrams illustrate phylogenetic incongruence in Betacoronaviruses. The phylogenetic tree built from the RdRp of Betacoronaviruses is on the left and the phylogenetic tree built from the S, E, M, and N genes is on the right for each tanglegram plot. A line connects genes from the same species. Species from the same subgenus are colored the same. Red: Sarbecovirus; orange: Hibecovirus; purple: Nobecovirus; green: Merbecovirus; blue: Embecovirus. **B** and **C.** Simplot analysis performed with Kimura (2-parameter), a window size of 1000 bp and a step size of 50 bp. The query genome for panel B is SARS-CoV-2 (accession number: NC_045512) and the query genome for panel C is Bat SL-CoVZC45 (accession number: MG772933). Recombinant regions detected with RDP4 are highlighted light grey.

### Analyzing recombination in SARS-CoV-2 and its close relatives

Due to the critical importance of understanding SARS-CoV-2 genome evolution, we performed a focused analysis on recombination in SARS-CoV-2 and its close relatives with SIMPLOT(Lole et al., 1999). The sequence identity of the bat coronavirus RATG13, is higher to SARS-CoV-2 than to the pangelin-derived coronaviruses MP789 and PCoV-GX_PL5 across the entire SARS-CoV-2 genome. The bat-derived coronavirus RmYN02 sequence identity is higher to SARS-CoV-2 than RATG13 over the majority of its length except for the region around S and ORF8 where the identity is lower. In fact, the sequence identity of RmYN02 across S and ORF8 is even lower than found in the pangolin coronaviruses MP789 and PCoV-GX_PL5. The decreased sequence identity of RmYN02 across S and ORF8 suggests recombination in these regions (relative to NC_045512:21259-24264 and NC_045512:27882-28259) (Figure 3B). The coronavirus CoVZC45, which was isolated from a bat in 2017, has a higher sequence identity to SARS-CoV-2 across its genome compared to SARS, except in the region corresponding to the majority of ORF1b (relative to NC_045512:11726-20372) (Figure 3C). These results suggest recombination occurred between the ancestors of SARS and SARS-CoV-2, implying the potential for recombination between SARS and SARS-CoV-2.

### TRS-B CS are more likely to be located in a breakpoint hotspot than in a random position in the genome

After locating TRS-B sites with SuPER, and systematically identifying recombination with RDP4, we investigated whether recombination is common at TRS-B sites. We determined whether TRS-B sites are located within a breakpoint hotspot, which is defined here as a region with breakpoint frequency higher than the 99th percentile of expected recombination breakpoint clustering assuming random recombination. We identified TRS-B sites from five subgenera located within recombination hotspots (Figure 4 and SFigure 2). In total, 8 out of 91 identified TRS-B sites were located within breakpoint hotspots. The recombination hotspots observed are more frequently co-located with TRS-B sites than expected (2-sample test for equality of proportions without continuity correction: p=2.2e-07). Interestingly, four of the eight TRS-B sites located within breakpoint hotspots proceed ORF S, an important determinant of host range. Two of the eight TRS-B sites located within breakpoint hotspots are found within Sarbecoviruses, one before ORF8 and one before gene N.

**Figure 4:**
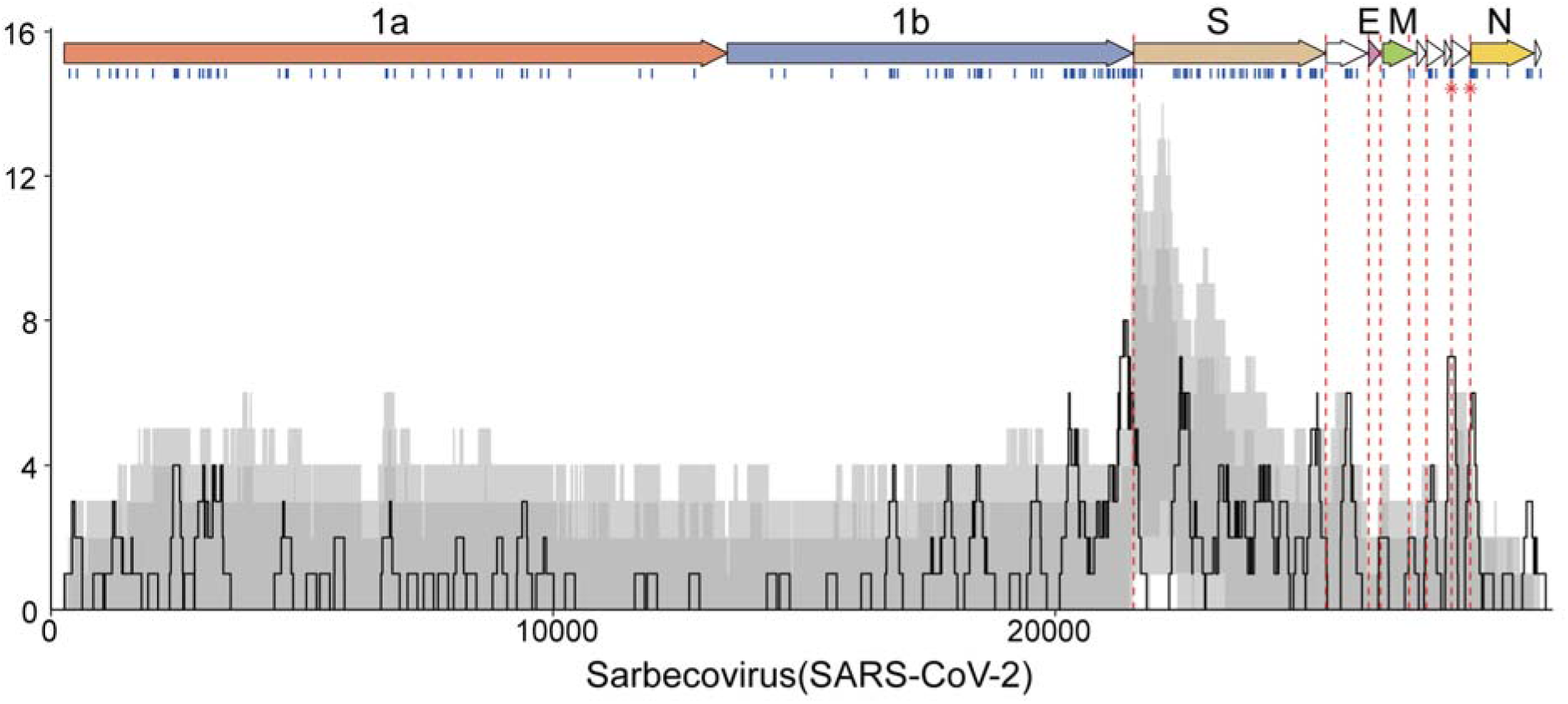
Recombination breakpoint density plot in Sarbecoviruses. Recombination breakpoint density plot illustrating breakpoint positions detected across 123 detectable events in Sarbecovirus using SARS-CoV-2 as the reference. Genes were plotted at the top of the panel as arrows. All detectable unique breakpoint positions are indicated by small vertical blue lines at the top of the graph. The number of breakpoints detected within the 200-bp region was counted and plotted as a solid black line. Light and dark grey areas respectively indicate the 95% and 99% intervals of expected breakpoint events assuming random recombination. Dashed red lines indicate the positions of TRS-B and red stars plotted at the red line to indicate that the TRS-B are located within breakpoint hotspots.

## DISCUSSION

Here, we sought to test the hypothesis that template switching at TRS-B sites during genome replication contributes to coronavirus genome recombination. SuPER, the tool we developed for identifying TRS sites in coronavirus genomes, will help to unravel this hypothesis. It should be noted that SuPER performs best given high-coverage RNA-seq data. While it still can predict TRS sites without RNA-seq, its performance declines and there are some instances, such as in Pedacovirus, where TRS sites cannot be reliably identified.

While the contribution of genome recombination generated at TRS-B sites can be difficult to dissect, we found that at 8 of 91 of TRS-B sites are located in breakpoint hotspots. However, this is likely an underestimate because our analysis is dependent on the number and diversity of available coronavirus genomes, and as of now, many coronavirus subgenera have only a handful of genomes available. We must also take into account the role of selection, in that only recombination events that produce viable viral genomes will survive and therefore be sequenced. For example, TRS-B sites tend to be intergenic, so recombination events that happen there are more likely to be viable, which could be an alternate explanation to the pattern we observe. As more coronavirus genomes are sequenced in the wake of the COVID-19 pandemic, the contribution of TRS-B sites to coronavirus genome recombination can be revisited. Experiments that examine recombination due to template switching at TRS-B sites that lessen the influence of selection pressure, such as direct RNA sequencing with Oxford Nanopore on cells co-infected with two coronaviruses, could further test this hypothesis. Overall, these results support but do not definitively confirm that recombinations at TRS-B due to template switching contribute to coronavirus genome recombination.

The worldwide pandemic caused by SARS-CoV-2 will likely lead to a global reservoir of SARS-CoV-2, as transmission to animals such as cats(Halfmann et al., 2020) and minks(Oreshkova et al., 2020) has been documented. A global reservoir of SARS-CoV-2 dramatically increases the chances that SARS-CoV-2 could recombine with other coronaviruses leading to recombinant viruses to which SARS-CoV-2 vaccines do not confer immunity. Furthermore, attenuated coronaviruses used for vaccines could revert to a virulent phenotype through recombination(Almazán et al., 2013; Graham et al., 2018; Pascual-Iglesias et al., 2019).

Therefore, it is of critical importance to understand factors and mechanisms that contribute to and underlie coronavirus genome recombination. Our results contribute to this through the systematic annotation of TRS sites in coronaviruses, which helps to inform which coronaviruses are at the highest risk of recombining with SARS-CoV-2 through TRS-B mediated template switching. Though we are limited by the number of available coronavirus genomes, we did not observe evidence for recombination between species in the subgenus Sarbecovirus and species in other Betacoronavirus subgenera or Alpha, Gamma, or Deltacoronaviruses. These results support that recombination between SARS-CoV-2 and other Sarbecoviruses is more likely and therefore, Sarbecoviruses should be the subject of intense surveillance. Overall, this work helps to inform and predict coronavirus genome recombination and could play a small but important role in preventing future coronavirus outbreaks.

## METHODS

### Identifying non-redundant coronavirus genomes

We downloaded 5517 complete genomes from the family of Coronaviridae from NCBI Virus (2020-05-19) and added one additional genome (EPI_ISL_412977 RmYN02) (Zhou et al., 2020) from GISAID. To remove the redundancy of the genomes, we used Mash(Ondov et al., 2016) to compute pairwise distance between the genomes (-d 0.01) and cluster them with MCL. We identified 506 clusters and selected one genome to represent each cluster(Shu and McCauley, 2017).

### Phylogenetic analysis

Protein sequences of RdRp, S, M, E, and N genes were retrieved and from each of the 506 representative genomes. Multiple sequence alignment was performed using Muscle(Edgar, 2004) with default parameters and phylogenetic trees built by FastTree from the alignment(Price et al., 2009). The phylogenetic tree based on RdRp was built to assign the taxonomy for those genomes without genus and subgenus assignments in NCBI Virus.

### Putative recombination events detection and analysis

Representative genomes from the same subgenera were aligned with Muscle(Edgar, 2004) and then used for RDP4(Martin et al., 2015) to detect recombination events. The GENECONV, MAXCHI, CHIMAERA, BOOTSCAN, SISCAN and 3SEQ methods implemented in the RDP4 package were used. Default RDP4 settings were used throughout the analysis. Events detected by four or more of the above methods were accepted and reported. Recombination density plots and hot-cold spots were identified using the RDP4 package. SIMPLOT(Lole et al., 1999) was employed to manually detect recombination events in the Sarbecovirus using a “query vs reference sequence” approach.

### Subgenomic mRNA Position Exploration with RNA-seq (SuPER)

SuPER (Subgenomic mRNA Position Exploration by RNA-seq) is designed to detect TRS sites in coronavirus genomes allowing for the delineation of sgmRNAs start sites. SuPER can be downloaded from: https://github.com/YiyanYang0728/SuPER. SuPER can use RNA-seq data to precisely delineate sgmRNA start sites and in the absence of RNA-seq, it uses the TRS-L site to predict TRS-B sites. The workflow of SuPER is divided into seven steps: 1) infer TRS-L in the 5’ UTR of reference genomes; 2) find split reads in the alignment of the RNA-seq to the coronavirus genome; 3) detect split sites supported by split reads as potential sgmRNA start sites; 4) refine assigned positions by identifying TRS-B in the reference genome; 5) associate the refined positions with possible downstream ORFs (<100 bp) if the genome annotation file is provided; 6) reconstruct site-specific 5’ end sgmRNA consensus sequences with split reads; 7) Report the alignment of TRS-L, TRS-B and the 5’ sgmRNA sequences.

#### Detection of TRS-L in reference genomes

The curated Stockholm file containing 5’ UTR alignment and consensus RNA secondary structure of major genera of Coronaviridae (namely Alphacoronavirus, Betacoronavirus, Gammacoronavirus, Deltacoronavirus) was downloaded from the Rfam database (http://rfam.xfam.org/covid-19), from which the CM files were generated by Infernal(Nawrocki and Eddy, 2013) with commands “cmbuild” and “cmcalibrate”. Given a reference genome and genus label, the first 150 bp of the genome was aligned using the corresponding genus CM file with command “cmalign” in Infernal. According to the consensus motif previously marked between SL2 and SL4 (often on SL3 if available) in the secondary structure, the counterpart sequence in the genome was eventually determined as its TRS-L.

#### Identifying sgmRNA start sites using RNA-seq

SuPER can use RNA-seq data to precisely detect TRS-B sites, which occur at the beginning of a sgmRNA. If the sequence mapping was obtained by a program using a similar algorithm to BWA(Li and Durbin, 2009), SuPER will find the split reads mapped both onto the leader sequence (within the first 10-150 bp in genome) and the sgmRNA 5’ end (between the coordinates 20000 and the end of the genome) on the same strand. On the other hand, if the sequence alignment is derived from reads mapped by a program considering RNA splicing such as HISAT2(Kim et al., 2015), SuPER will use junction reads instead. The split or junction reads then define a putative sgmRNA start site. The putative sgmRNA start site is further refined by searching for the TRS-B sequence. The region spanning 30bp upstream and downstream of the putative sgmRNA start site is searched with a sliding window with the same length as TRS-L. The sequence with minimal hamming distance from TRS-L is assigned as the TRS-B site. In addition, if a genome annotation file is provided, SuPER will try to connect the sgmRNA start site to the nearest downstream ORF within 170 bp.

#### Identifying sgmRNA start sites without RNA-seq

Although the best results are obtained when using RNA-seq, SuPER still functions when only a reference genome and annotation are provided. After inferring the TRS-L site in the reference genome, SuPER is capable of finding possible TRS-B sites throughout the whole genome with a hamming distance from TRS-L less than 1 or less than 2 if a downstream ORF exists. In the situation of multiple positions associated with the same ORF, the position with minimal hamming distance and closest to the ORF is assigned as the TRS-B site.

## ACKNOWLEDGMENTS

Y.Y., W.Y., X.J. were supported by the Intramural Research Program of the NIH, National Library of Medicine. A.B.H. is supported by the University of Maryland Department of Cell Biology and Molecular Genetics and the Center for Bioinformatics and Computational Biology.

## COMPETING INTERESTS

The authors declare no competing interests.

## Supplementary information

### Supplementary Tables

Supplementary Table 1: Representative genomes for coronaviruses and host range

Supplementary Table 2: Coronavirus species with RNA-seq data available

Supplementary Table 3: Identification of TRS-L/B for 506 representative genomes

Supplementary Table 4: SuPER benchmarks

### Supplementary Data

Supplementary Data 1: Recombination events detected by RDP4

Supplementary Data 2: Analysis of SuPER on coronavirus genomes with RNA-seq data

### Supplementary Figures

Supplementary Figure 1. The secondary structure of TRS-L

Supplementary Figure 2. Density plot of recombination in each subgenus

## Notes

### Competing Interest Statement

The authors have declared no competing interest.

