## Supplementary figures and images for "Characterizing transcriptional regulatory sequences in coronaviruses and their role in recombination"

### Supplementary Figures

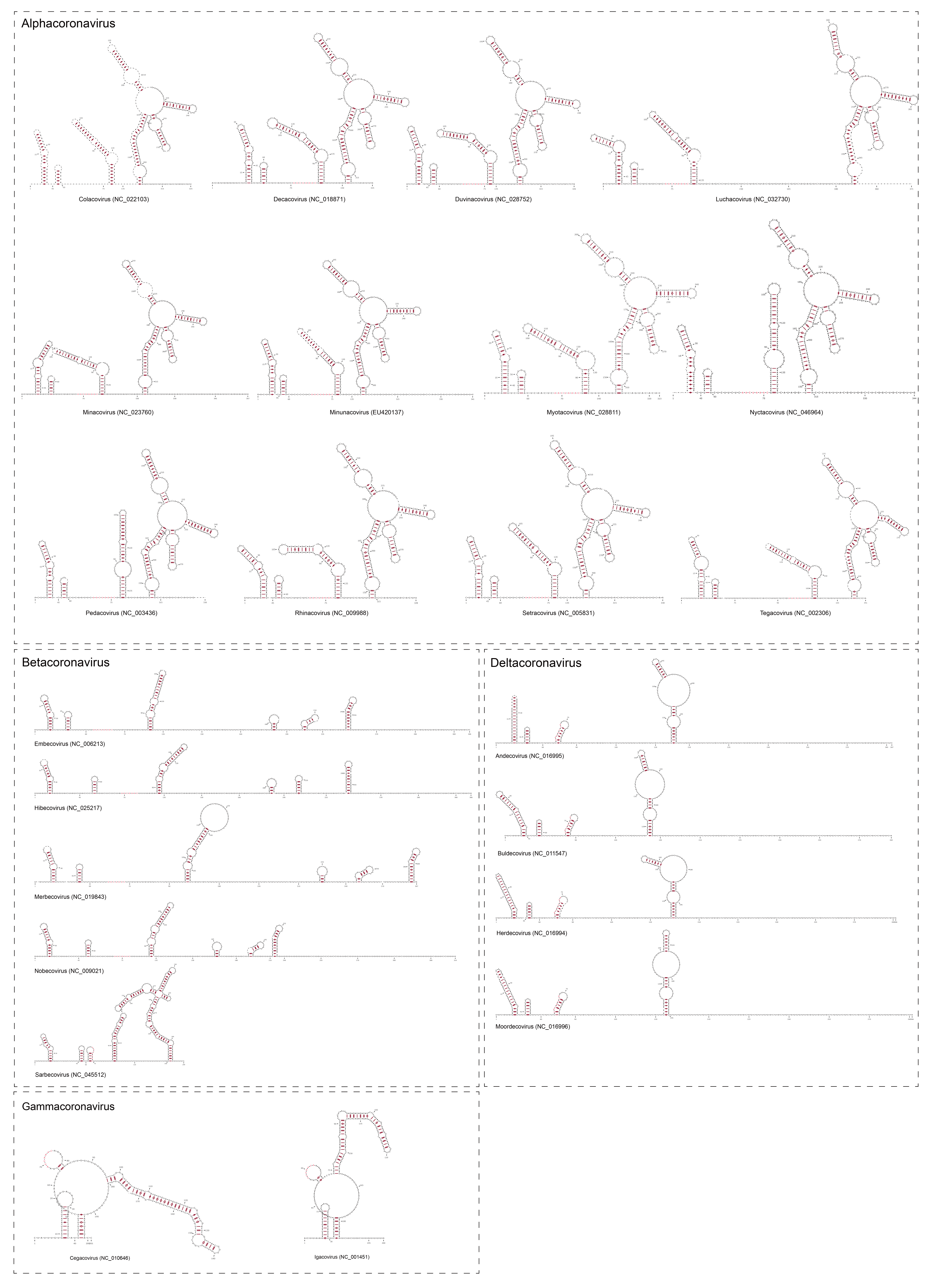

SFigure 1. The structure of TRS-L

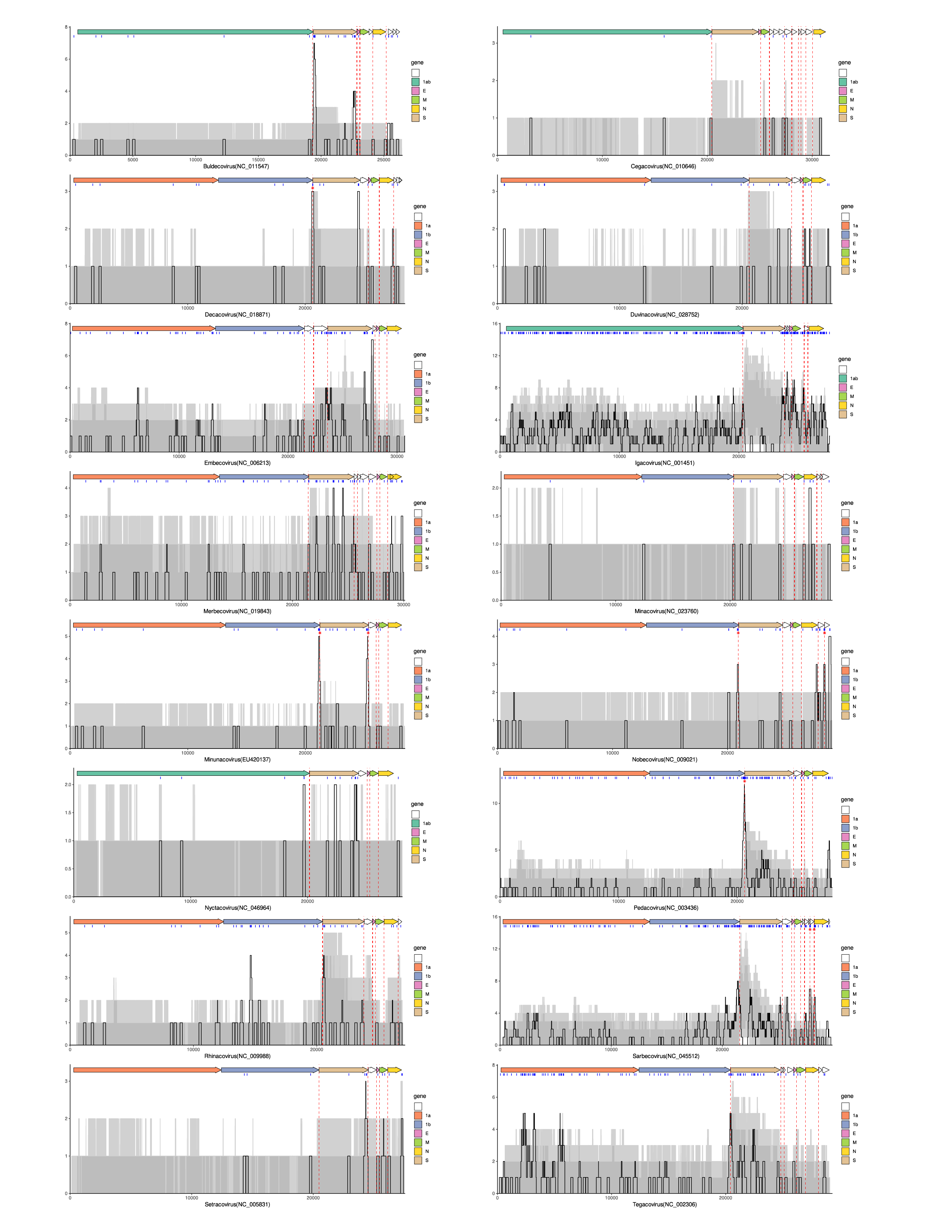

SFigure 2. The density plot of recombination in each subgenus
